# Patient-Derived hiPSC-Cardiomyocytes and Engineered Heart Tissues Reveal Distinct Functional Phenotypes in Inherited Cardiomyopathies

**DOI:** 10.64898/2026.08.28.747951

**Authors:** Saana Pohjavaara, Qasim A. Majid, Leo Huttunen, Katriina Aalto-Setälä, Heikki Ruskoaho, Mika J. Välimäki, Sini M. Kinnunen, Virpi Talman

**Affiliations:** Drug Research Program and Division of Pharmacology and Pharmacotherapy, Faculty of Pharmacy, University of Helsinki, Helsinki, Finland; Department of Pharmacology and Toxicology, Institute of Pharmaceutical Sciences, University of Graz, Graz, Austria; Faculty of Medicine and Health Technology and BioMediTech Institute, Tampere University, Tampere, Finland; Heart Hospital, Tampere University Hospital, Tampere, Finland; Research Centre for Integrative Physiology and Pharmacology, Institute of Biomedicine, Faculty of Medicine, University of Turku, Turku, Finland

**Keywords:** hypertrophic cardiomyopathy, dilated cardiomyopathy, engineered heart tissues, hiPSC-derived cardiomyocytes, GATA4

## Abstract

**Background:** Hypertrophic and dilated cardiomyopathies (HCM and DCM) are the most common inherited cardiomyopathies. However, genotype-specific molecular and functional cardiomyocyte phenotypes and responses to neurohormonal stimulation remain incompletely understood. Here, we investigated whether patient-derived HCM and DCM cardiomyocytes exhibit distinct baseline phenotypes or differential responses to hypertrophic stimulation and pharmacological treatment.

**Methods:** Three human-induced pluripotent stem cell (hiPSC) lines were used: a control line, an HCM patient-derived line carrying a *MYBPC3* mutation, and a DCM patient-derived line carrying an *LMNA* mutation. The cells were differentiated into hiPSC-cardiomyocytes, which were exposed to endothelin-1 and the GATA4-targeted compound 3i-1262, followed by transcriptional and protein expression analyses. In addition, engineered heart tissues (EHTs) were generated and cultured for 40 days, with endothelin-1 and 3i-1262 treatment applied during the final 20 days. Lastly, β-adrenergic stimulation with isoprenaline was performed. EHT contractile function was quantified using MUSCLEMOTION.

**Results:** Patient-derived hiPSC-cardiomyocytes exhibited genotype-dependent responses to endothelin-1 at the transcriptional and protein levels. DCM-cardiomyocytes failed to maintain structural integrity in the EHTs, resulting in tissue fracture or cessation of beating. Functional analyses demonstrated distinct baseline contractile properties between control and cardiomyopathy EHTs, as well as differential responses to endothelin-1 and isoprenaline.

**Conclusions:** Patient-derived hiPSC-cardiomyocytes exhibit genotype-specific molecular and functional phenotypes. EHTs generated from HCM hiPSC-derived cardiomyocytes showed a progressive decline in apparent force, whereas DCM EHTs fractured over time, suggesting mutation-associated phenotypes in 3D cardiac tissue models. These findings highlight the utility of hiPSC-based cardiac models for investigating molecular and functional disease mechanisms and pharmacological responses in inherited cardiomyopathies.

## Introduction

Hypertrophic and dilated cardiomyopathies (HCM and DCM) are among the most common inherited cardiac disorders, affecting approximately 1 in 500 and 1 in 250 individuals globally, respectively ^1^. These disorders are characterized by profound structural and functional abnormalities of the heart and are associated with major adverse outcomes, including heart failure, arrhythmias, and sudden cardiac death ^2–4^. Advances in identifying disease-causing genetic variants and elucidating the genetic basis of cardiomyopathies have further highlighted their substantial contribution to cardiovascular morbidity and mortality ^1^.

Transgenic animal models and derived in vitro systems have provided important insights into the mechanisms underlying cardiomyopathies ^5,6^. However, their translational relevance is limited by interspecies differences in cardiac physiology, such as heart rate and electrophysiology, gene and protein expression, including myosin heavy chain isoform profiles, and the limited availability and short lifespan of primary rodent cardiomyocytes. Human induced pluripotent stem cells (hiPSCs) provide a renewable and patient-specific platform for modeling cardiac diseases ^7^. hiPSC-derived cardiomyocytes (hiPSC-cardiomyocytes) carrying disease-associated variants can recapitulate key features of HCM and DCM, including altered gene expression, contractility, cellular architecture, and stress responses, thereby allowing detailed investigation of genotype-specific phenotypes in a human context ^7–12^. Among disease-causing variants, the *MYBPC3* (myosin binding protein C3) Gln1061X mutation, associated with HCM, leads to sarcomeric disorganization and impaired contractile function, whereas the *LMNA* (lamin A/C) S143P mutation, associated with DCM, disrupts nuclear envelope integrity and impairs mechanotransduction ^8,9,13^.

Advanced tissue engineering approaches, such as three-dimensional engineered heart tissue (EHT), promote the structural and functional maturation of hiPSC-cardiomyocytes and enable quantitative assessment of apparent contractile force and kinetics, providing physiologically relevant readouts of disease phenotypes ^14–16^. These constructs recapitulate key pathological features of cardiomyopathies beyond those observed in traditional monolayer cultures, including impaired force generation, altered passive mechanical properties, altered contraction kinetics, and changes in sarcomeric protein composition^17,18^.

Beyond basal characterization of HCM and DCM hiPSC-cardiomyocytes, it is essential to understand how cardiomyocytes respond to external stimuli. While basal conditions reveal mutation-driven differences, they do not capture how cardiomyocytes respond to pathological stress. Endothelin-1 is a well-established pro-hypertrophic peptide that induces changes in gene and protein expression, as well as cellular morphology, in cardiomyocytes ^19–23^. Endothelin-1-induced hypertrophic responses are mediated in part by the transcription factor GATA4, whose phosphorylation and increased DNA-binding activity promote the expression of hypertrophy-associated genes such as *NPPA* and *NPPB*, which encode atrial and B-type natriuretic peptides, respectively ^24–27^. The GATA4-targeted compound 3i-1262 has been shown to attenuate endothelin-1-induced upregulation of hypertrophic genes in hiPSC-cardiomyocytes ^28^. Mutation-specific responses to endothelin-1 may reveal functional and molecular differences that are not apparent under basal conditions. Therefore, in this study, we investigated how the *LMNA* S143P and *MYBPC3* Gln1061X mutations influence endothelin-1-induced molecular and functional responses and assessed whether GATA4-targeted compound 3i-1262 could modulate these responses. In addition, patient-derived EHTs were generated to assess contractile function in untreated and endothelin-1 or 3i-1262-treated cardiomyocytes.

## Methods

### Human-induced pluripotent stem cell-derived cell lines

The control cell line iPS(IMR90)-4 was purchased from WiCell (Madison, WI, USA)^29^. Generation and characterization of the patient-derived hiPSC lines have been described earlier ^8,9^. Cardiomyopathy-related lines were purchased from the iPS Cells core facility of the University of Tampere. DCM-related hiPSCs carry a p.S143P mutation in the *LMNA,* while the HCM-related hiPSCs carry a Gln1061X mutation in the *MYBPC3*. The presence of p.S143P (TCC to CCC) in the *LMNA* gene in the DCM hiPSCs and Gln1061X truncating mutation (CAG to TAG) in the *MYBPC3* gene in the HCM hiPSCs has been confirmed previously with DNA sequencing ^30^.

### Cell culture and differentiation into hiPSC-derived cardiomyocytes

All reagents used in the cell culture were obtained from Gibco, unless otherwise specified. Cell culture and hiPSC differentiation have been described previously ^31^. Briefly, hiPSCs were cultured in feeder-free conditions on Matrigel-coated plates using Essential 8™ medium at 37 °C in a humidified 5% CO_2_ atmosphere. Cells were passaged approximately every four days at a 1:15 ratio using Versene, with ROCK inhibitor Y-27632 (Tocris; 10 µM) added for 24 h following each passage. Cardiomyocyte differentiation was initiated at 70–80% confluency (day 0) using 6–8 µM CHIR99021 (Tocris) in RPMI 1640 supplemented with B-27 Minus Insulin (RB-) ^32^. On day 1–2, cells were refreshed with RB-without CHIR99021, followed by addition of RB-containing the Wnt inhibitor Wnt-C59 (Tocris; 2.5 µM) on day 3. Media changes with fresh RB- were performed on days 5, 7, and 9. To purify the culture, metabolic selection was carried out on days 11 and 13 by culturing cells in glucose-free RPMI with B-27. From day 15 onwards, cardiomyocytes were maintained in RPMI 1640 supplemented with B-27 (RB+) and fed twice weekly.

For downstream experiments, cells were dissociated on days 15–18 using TrypLE™ Select following the manufacturer’s protocol. Cardiomyocytes were seeded onto Matrigel-coated plates and cultured in RB+ supplemented with 10% heat-inactivated fetal bovine serum (FBS) and the ROCK inhibitor Y-27632. Cells were plated as follows: for high-content analysis (HCA), 1–1.3x10^4^ cells per well in black, clear-bottomed PhenoPlate 96-well plates (Revvity); for gene expression analysis, 5x10^5^cells per well in standard 12-well plates; for western blotting, 1x10^6^ cells per well in standard 6-well plates; and for engineered heart tissue (EHT) generation, 1x10^6^cells per well in standard 6-well plates. After 48 h, the medium was replaced with fresh RB+, and cultures were maintained until hypertrophy induction. Prior to experimentation, cultures were visually inspected to confirm spontaneous beating, and experiments were only performed if clear contractions were observed. For immunofluorescence analyses, downstream assays were conducted only if more than 80% of cells were confirmed to be cardiomyocytes by cardiac troponin T (cTnT) staining.

### Hypertrophy induction and 3i-1262 treatment for immunofluorescence staining, qPCR and Western blot

Days 29–41 cells were used for immunofluorescence staining and high-content analysis, as well as gene and protein expression studies. For this, cells were treated with the GATA4-targeted compound 3i-1262 (30 µM)^28^ or DMSO vehicle (0.1%) for 1 h prior to a 24-hour exposure to endothelin-1 (100 nM). The schematic for the experimental design is presented in Figure 1A.

**Figure 1.**
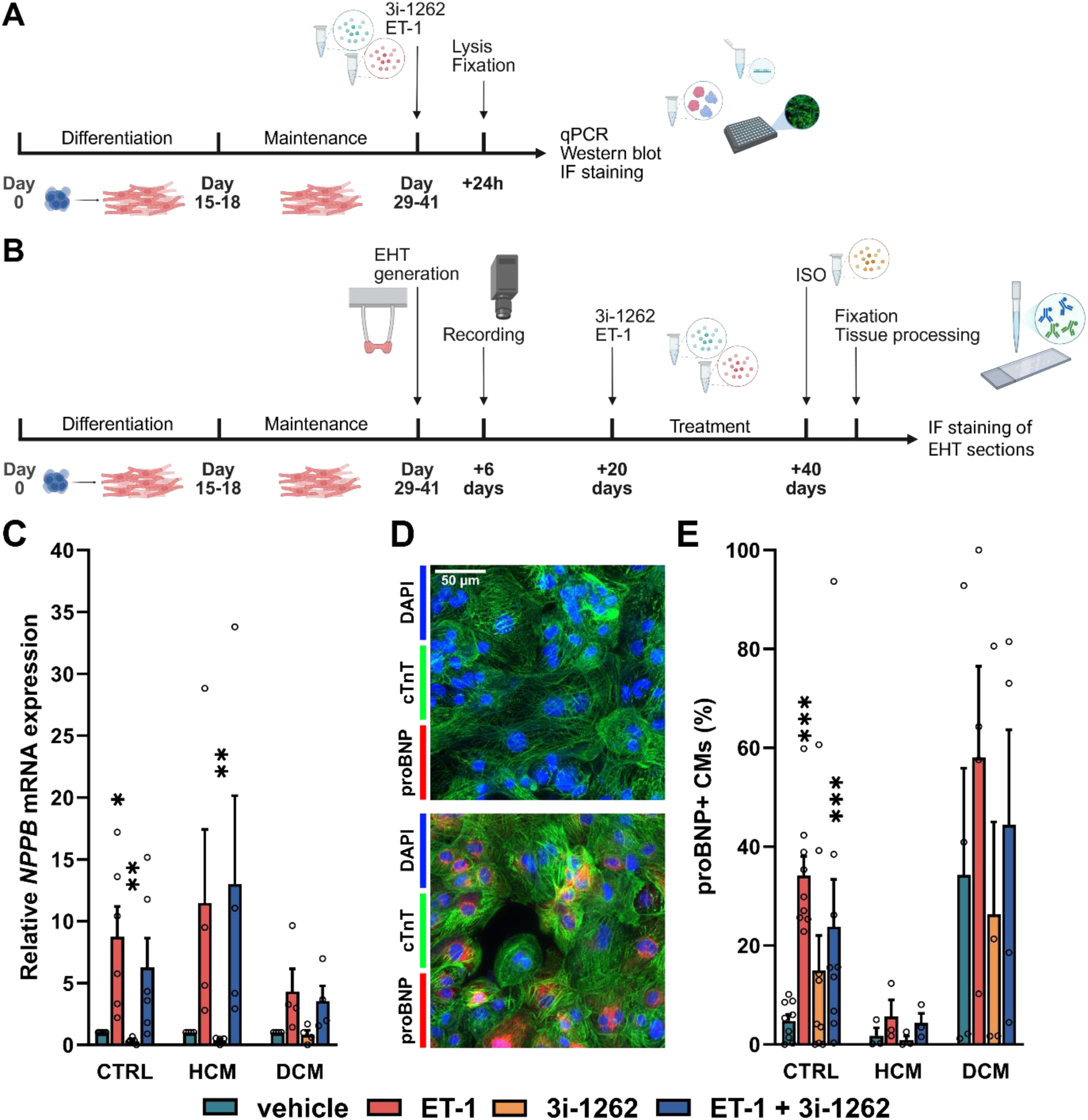
Endothelin-1-induced *NPPB* and proBNP upregulation in hiPSC-cardiomyocytes. A) Schematic overview of the experimental design for qPCR, Western blot, and immunofluorescence staining. B) Schematic overview of the experimental design for EHT generation, recording, and immunofluorescence staining of the EHT sections. C) Relative *NPPB* expression in hiPSC-cardiomyocytes in response to pro-hypertrophic endothelin-1 stimuli and 3i-1262 treatment. Results are presented as mean + SEM and normalized to the respective control sample. D) Representative image of control hiPSC-cardiomyocytes treated with vehicle or endothelin-1. Cells were stained for DAPI (blue), cardiac troponin T (cTnT; green) and pro-B type natriuretic peptide (proBNP; red). E) Percentage of proBNP+ cardiomyocytes based on the average intensity of proBNP in the perinuclear area of hiPSC-cardiomyocytes. Results are presented as mean + SEM. n = 3–9, *p<0.05; **p<0.01; ***p<0.001 compared to the respective vehicle sample (independent samples t-test or Welch’s t-test, as appropriate). ET-1, endothelin-1, IF, immunofluorescence; EHT, engineered heart tissue; ISO, isoprenaline; CTRL, control cardiomyocytes; HCM, HCM cardiomyocytes; DCM, DCM cardiomyocytes. Schematics in A and B were created with Biorender.com.

### Immunofluorescence staining and high-content analysis

To enhance pro-B-type natriuretic peptide (proBNP) detection, Brefeldin A (Invitrogen) was added 3 hours before fixation. The cells were washed twice with phosphate-buffered saline (PBS), fixed with 4% paraformaldehyde at room temperature for 15 min, and permeabilized with 0.1% Triton X-100 (AppliChem) in PBS for 10 min. Non-specific binding sites were blocked with 4% FBS in PBS for 45 min. Thereafter, primary antibodies were diluted in 4% FBS in PBS at 1:800 and incubated for 60 min at room temperature: cardiac troponin T (ab 45932, Abcam) and proBNP (ab11315, Abcam). Cells were then washed 3x5min with PBS and Alexa Fluor ® -conjugated secondary antibodies (Invitrogen) at 1:200 and 4′,6-diamidino-2-phenylindole (DAPI; Sigma-Aldrich) at 1µg/mL were incubated for 45 min at room temperature, protected from light. Cells were washed with 3x5 min PBS and stored at 4°C in PBS, protected from light until imaged. ImageXpress Micro Confocal high-content screening system (Molecular Devices) with a Nikon 10x Plan Apo Lambda air objective was used to image the cells. MetaXpress software (Molecular Devices) was used to analyze proBNP intensity, as described earlier ^31^. Furthermore, the proportion of cardiomyocytes, as determined by DAPI and cTnT staining, was quantified in each culture, and only cultures with ≥80% cardiomyocytes were included in downstream analyses.

### RNA isolation, cDNA synthesis, quantitative PCR

Total RNA was isolated with NucleoSpin RNA kit (Macherey–Nagel) following the manufacturer’s instructions. RNA concentration and quality were analyzed with a NanoDrop Microvolume Spectrophotometer (Thermo Fisher Scientific). Then, 115– 1047 ng of RNA was reverse-transcribed to single-stranded cDNA in 10–20 µl reactions with Transcriptor First Strand cDNA Synthesis Kit (Roche) using random hexamer primers and MJ Mini Personal thermal cycler (Bio-Rad). Equal amounts of RNA from both the cardiomyopathy cell line and the corresponding control were reverse-transcribed within each biological replicate (n = 1).

The resulting cDNA was diluted 1:10 in nuclease-free water and stored at −20 °C. Gene expression was quantified using commercial TaqMan® Gene Expression Assays (Thermo Fisher Scientific; Supplementary table 1) with the LightCycler® 480 ProbesMaster reagent (Roche) according to the manufacturer’s instructions. Reactions were performed in a 10 µl total volume, including 4.5 µl of diluted cDNA, in white 384-well LightCycler® 480 plates and run on a LightCycler® 480 Real-Time PCR System (Roche). Each reaction was carried out in triplicate, and Grubbs’ test (α = 0.05) was applied to identify and exclude any outliers among technical replicates. Relative gene expression was calculated using the 2^−ΔΔCt method, with *ACTB* and *18S* rRNA serving as endogenous controls ^31^.

### Western blot

Brefeldin A (Invitrogen) was added to the cells 3 hours before cell lysis. Cells were lysed in 1% sodium dodecyl sulfate in 50 mM Tris-HCl (pH 7.5). Total protein concentrations were determined using the Pierce™ BCA Protein Assay Kit (Thermo Scientific). Total protein (15 µg) was loaded into 10% Mini-PROTEAN® TGX Stain-Free™ Protein gels (Bio-Rad). Control hiPSC-cardiomyocyte samples treated with endothelin-1 from an independent differentiation were included as positive controls and loaded on the gel alongside the samples of interest ^28^. Gels were exposed to UV light for 5 min to activate trihalo compound-protein interactions, after which proteins were transferred to nitrocellulose membranes using the Trans-Blot® Turbo™ system (Bio-Rad).

Non-specific binding was blocked with 5% non-fat dry milk in Tris-buffered saline containing 0.1% Tween 20 (TTBS) for 1 h at room temperature. Membranes were incubated overnight at 4 °C with primary antibody for proBNP (Abcam, ab13115) at 1:1000 in 5% milk–TTBS. Following primary incubation, membranes were treated for 1 h with 1:2000 HRP-conjugated secondary antibody (anti-mouse, Cell Signaling Technology, 7076).

Protein bands were visualized using enhanced chemiluminescent substrate SuperSignal™ West Femto Maximum Sensitivity (Thermo Scientific), with imaging performed on a ChemiDoc™ MP System (Bio-Rad). Band intensities were quantified using Fiji ImageJ 1.53 software and normalized to the total protein content of each sample.

### Generation of engineered heart tissues

Engineered heart tissues (EHT) were fabricated as described earlier^33,34^. Briefly, hiPSC-cardiomyocytes between days 29 and 41 of differentiation were dissociated by incubating them for 3–4 hours in a dissociation media comprised of Hanks’ Balanced Salt Solution containing collagenase type 2 (200 U/mL, 156 µl/cm²) and 10 µM ROCK inhibitor Y-27632 under standard culture conditions. The dissociation agent was neutralized with a neutralization media (RB+, 10% FBS, and 12 µg/mL DNase 1 (Applichem)), and cells were centrifuged at 100g for 15 min. Cells (1x10^6^ cells/EHT) were then resuspended in a reconstitution mixture of non-cardiomyocyte media (NKM), 2X Dulbeccoʹs Modified Eagle Medium, bovine fibrinogen at 200 mg/mL, and bovine thrombin 100 units/mL. Casting molds were prepared by placing the Teflon spacer in a standard 24-well plate and adding 2 mL of 2 % UltraPure™ Agarose (Sigma Aldrich) in PBS per well. The spacer was removed following agarose solidification, and silicon posts (DiNABIOS) were placed into the wells. The reconstitution mixture was then dispensed into the molds with spacers and incubated for 90 min in standard culture conditions, whereafter 300 µl of NKM media was added to the wells to rehydrate the EHTs. After 30 min, the produced EHTs were moved to the adjacent wells containing 1.5 mL of EHT media (10% FBS, 10 µg/mL insulin, 33 µg/mL bovine aprotinin in Knockout DMEM). EHTs were fed with warm EHT media every other day 30 min before video recordings. The schematic for the experimental design is shown in Figure 1B.

### EHT recording and contractile analysis

EHT recording was initiated on day 6 post-production, as the embedded cardiomyocytes typically require time to achieve synchronous contraction and generate measurable force^34^. Recordings of the total EHT area were subsequently obtained every other day until day 40 using a Basler ace acA1440-220uc camera controlled via the Basler pylon Viewer software. Each recording comprised 1,000 frames acquired over a 10-second interval. Contraction kinetics were then analyzed with a MUSCLEMOTION plugin in ImageJ ^15^. To calculate the apparent generated force, the image scale (1231.83 pixels/cm) was determined in ImageJ. The total EHT length (L_EHT_) was measured in a relaxed state, while the interpost distance was measured in both the relaxed and contracted states. Fractional shortening (FS) was calculated from the change in interpost distance and expressed as a percentage. Post deflection (δ, mm) was then calculated as follows:

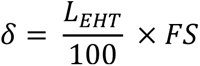

Post deflection was subsequently converted from millimeters to micrometers. The apparent generated force (mN) was calculated based on Vandenburgh et al. (2008)^16^ using the following formula:

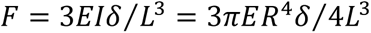

where *E* is Young’s modulus (MPa), *I* is the moment of inertia (mm^4^), *L* is the length of the silicon post (mm), *R* is the radius of the post (mm), and δ is the post deflection (µm).

### Hypertrophy induction, 3i-1262 treatment and β-adrenergic stimulation for EHTs

Four EHTs were produced from each differentiation to have four treatment groups: endothelin-1 (100nM), 3i-1262 (30 µM), their combination and vehicle control (DMSO 0.1%). Treatments were initiated on day 20 post-production and readministered at every feeding until day 40. After the final treatment and recordings, EHTs were exposed to 1 µM isoprenaline^14^ to assess the effects of β-adrenergic stimulation on contraction kinetics, with recordings taken at 5, 10, 30, and 60 minutes post-exposure.

### Immunofluorescent staining of paraffin-embedded EHTs

After 60 minutes of incubation with isoprenaline, EHTs were washed twice with PBS, fixed with 4% PFA overnight at 4°C, and stored in PBS until tissue processing. EHTs were embedded in paraffin and sectioned longitudinally to produce 4 µm sections. Thereafter, sections were deparaffinized using a series of xylene and ethanol washing steps, and underwent antigen retrieval in heated sodium citrate buffer, followed by permeabilization with 0.2% Triton X-100 in Tris-buffered saline (TBS) for 10 minutes at room temperature. To block non-specific binding, the sections were incubated in 5% FBS in TTBS for 1 h at room temperature. Sections were then incubated in a primary antibody solution containing anti-cardiac troponin T (Abcam, ab45932) at 1:400 in blocking solution overnight at 4°C. Alexa Fluor® -conjugated secondary antibodies (Invitrogen) were added at 1:200 in blocking solution and incubated for 1 h at room temperature, and DAPI (Sigma-Aldrich) at 1µg/mL was incubated for 10 minutes at room temperature, protected from light. Sections were washed with 3x10 min TBS and mounted with coverslips. Sections were imaged with a Leica DMi8 widefield microscope with a 5x objective (N PLAN 5x/0.12, dry) and a 10x objective (N PLAN 10x/0.45, dry).

### Statistical analysis

For qPCR analyses, the mean of three technical replicates was used to represent one biological replicate (n = 1). Each treatment group consisted of 3-9 independent experiments derived from separate differentiations. qPCR and Western blot data are presented as mean ± standard error of the mean (SEM). Homogeneity of variance was assessed using Levene’s test; independent-samples t-tests were used when equal variances were assumed, whereas Welch’s t-test was used otherwise. Exact p-values are reported throughout.

EHT kinetic parameters were analyzed using linear mixed-effects models with group, treatment, day and their interactions included as fixed effects, and repeated measurements within each EHT accounted for by a random intercept. Fixed effects were evaluated using Type III tests. Post hoc pairwise comparisons were adjusted using the Bonferroni method to control the risk of type I error arising from multiple comparisons across group, treatment, and time. Effect estimates are reported with 95% confidence intervals and exact p-values in the Results section. The number of biological replicates is presented in Supplementary tables 2 – 4.

A priori power calculations were not performed because reliable estimates of effect sizes and variability were unavailable for the studied hiPSC-cardiomyocytes and EHT models. Sample sizes were based on the number of independent differentiations and EHTs that could be generated and analyzed using established protocols. All statistical analyses were performed using IBM SPSS Statistics version 31.0.0.

## Results

### Effects of endothelin-1 and the GATA4-targeted compound 3i-1262 on *NPPB* and proBNP expression

To assess endothelin-1-induced hypertrophic responses in cardiomyocytes, the expression levels of *NPPB* (encoding B-type natriuretic peptide) mRNA and proBNP protein were analyzed. There was no difference in the basal expression of *NPPB* across the cell lines (Supplementary figure 1). Endothelin-1 treatment for 24 hours increased the *NPPB* expression across all cell lines, with an 8.7-fold increase in control, 11.4-fold in HCM and 4.3-fold in DCM cardiomyocytes; however, this upregulation was not statistically significant in HCM or DCM cardiomyocytes (p = 0.025, p = 0.176 and p = 0.165, respectively) (Figure 1C). Treatment with compound 3i-1262 alone reduced *NPPB* expression by 50% in control cardiomyocytes, whereas no change was observed in HCM and DCM cardiomyocytes (Figure 1C). ProBNP expression was analyzed based on immunofluorescence staining, and proBNP intensity was quantified in the perinuclear area (Figure 1D – E). We observed an increase in proBNP expression in control cardiomyocytes (7.1-fold), whereas the increases in HCM (3.2-fold) and DCM (1.7-fold) cardiomyocytes were not statistically significant (p = 0.374 and p = 0.436, respectively). We also assessed proBNP expression in whole-cell lysates using Western blotting.

Endothelin-1 increased proBNP expression to 4.4-fold in control (p = 0.181, not significant) and 8.9-fold in HCM cardiomyocytes (p = 0.122, not significant) (Supplementary figure 3 4). In DCM cardiomyocytes, endothelin-1 did not increase proBNP expression. In control cardiomyocytes, 3i-1262 tended to attenuate the endothelin-1-induced increase in proBNP expression to 1.6-fold and to 4-fold in HCM cardiomyocytes; however, these effects were not significant (p = 0.227 and p = 0.270).

### Hypertrophy-associated gene expression in hiPSC-cardiomyocytes in response to endothelin-1 and the GATA4-targeted compound 3i-1262

We next analyzed other genes associated with the hypertrophic response (Figure 2). In basal conditions, the only significant difference between cell lines was higher expression of *ACTC1*, encoding α-cardiac actin, in DCM cardiomyocytes (Supplementary figure 1). After endothelin-1 treatment, the expression of *NPPA*, which encodes the natriuretic peptide, increased in control cardiomyocytes to 2.2-fold, with a similar, non-significant increase observed in HCM and DCM cardiomyocytes. The expression of *MYH6*, which encodes the myosin heavy chain α-isoform, was downregulated by 40% in control cardiomyocytes. Interestingly, endothelin-1 induced a marked increase in *ACTA* expression, which encodes the skeletal muscle α-actin, in control (57-fold), HCM (18-fold), and DCM (12-fold; p = 0.285, not significant) cardiomyocytes. The GATA4-targeted compound 3i-1262 alone did not alter the expression of these genes. When 3i-1262 was combined with endothelin-1 treatment, *ACTA1* and *TNNT2* expression increased in control cardiomyocytes, while *ACTA1* expression increased in HCM cardiomyocytes and *NPPA* expression increased in DCM cardiomyocytes when compared to untreated cells. However, in the case of *ACTA1*, the increase is most likely due to endothelin-1, rather than to the combined effect of the two.

**Figure 2.**
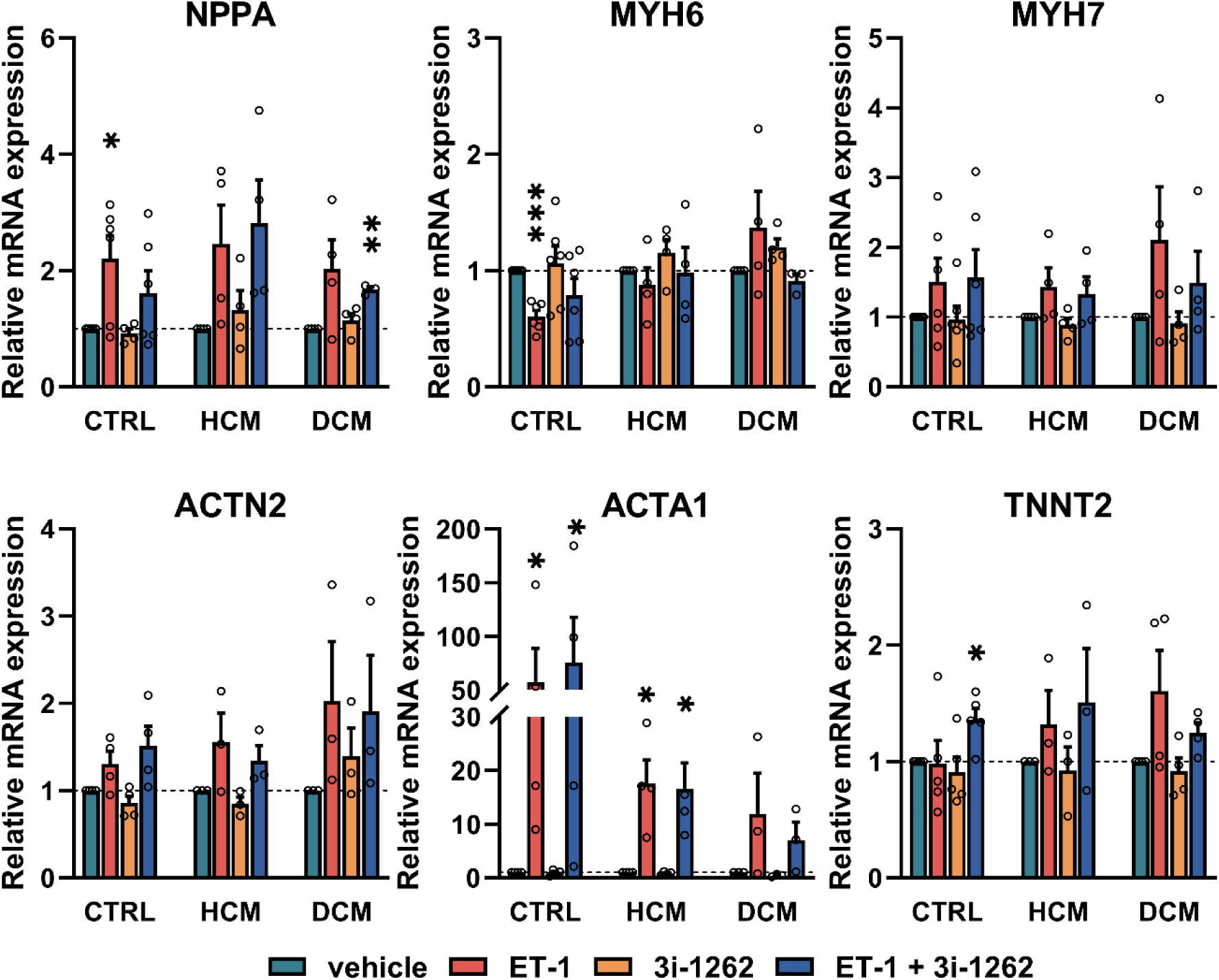
Gene expression of hypertrophy-related genes in hiPSC-cardiomyocytes after 24 hours of endothelin-1 exposure and 3i-1262 treatment. Results are presented as mean + SEM and normalized to the respective vehicle sample. n = 3–6, *p<0.05; **p<0.01 compared to the respective vehicle sample (independent samples t-test or Welch’s t-test, as appropriate). CTRL, control cardiomyocytes; HCM, HCM cardiomyocytes; DCM, DCM cardiomyocytes; ET-1, endothelin-1.

### Cardiomyocyte structural and regulatory genes

Next, we expanded the gene panel to include genes previously shown to be regulated by other hypertrophic stimuli, particularly mechanical stretching ^30,31^ (Figure 3). In control cardiomyocytes, *MYPBC3* mRNA expression was reduced following ET-1 administration, whereas galanin (*GAL*) and cysteine- and glycine-rich protein 3 (*CSRP3*) expression was increased. However, no effects were observed in the disease lines. Interestingly, endothelin-1 reduced *SLC16A9* (solute carrier family 16 member 9) expression in HCM cardiomyocytes, contrasting with the genotype-independent downregulation of *SLC16A9* previously observed in response to mechanical stretch ^30^. Consistent with previous findings, the GATA4-targeted compound 3i-1262 increased *SLC16A9* expression by 12-fold in control and 6.7-fold in HCM cardiomyocytes, with a similar, non-significant 5.1-fold trend in DCM cells (p = 0.142).

**Figure 3.**
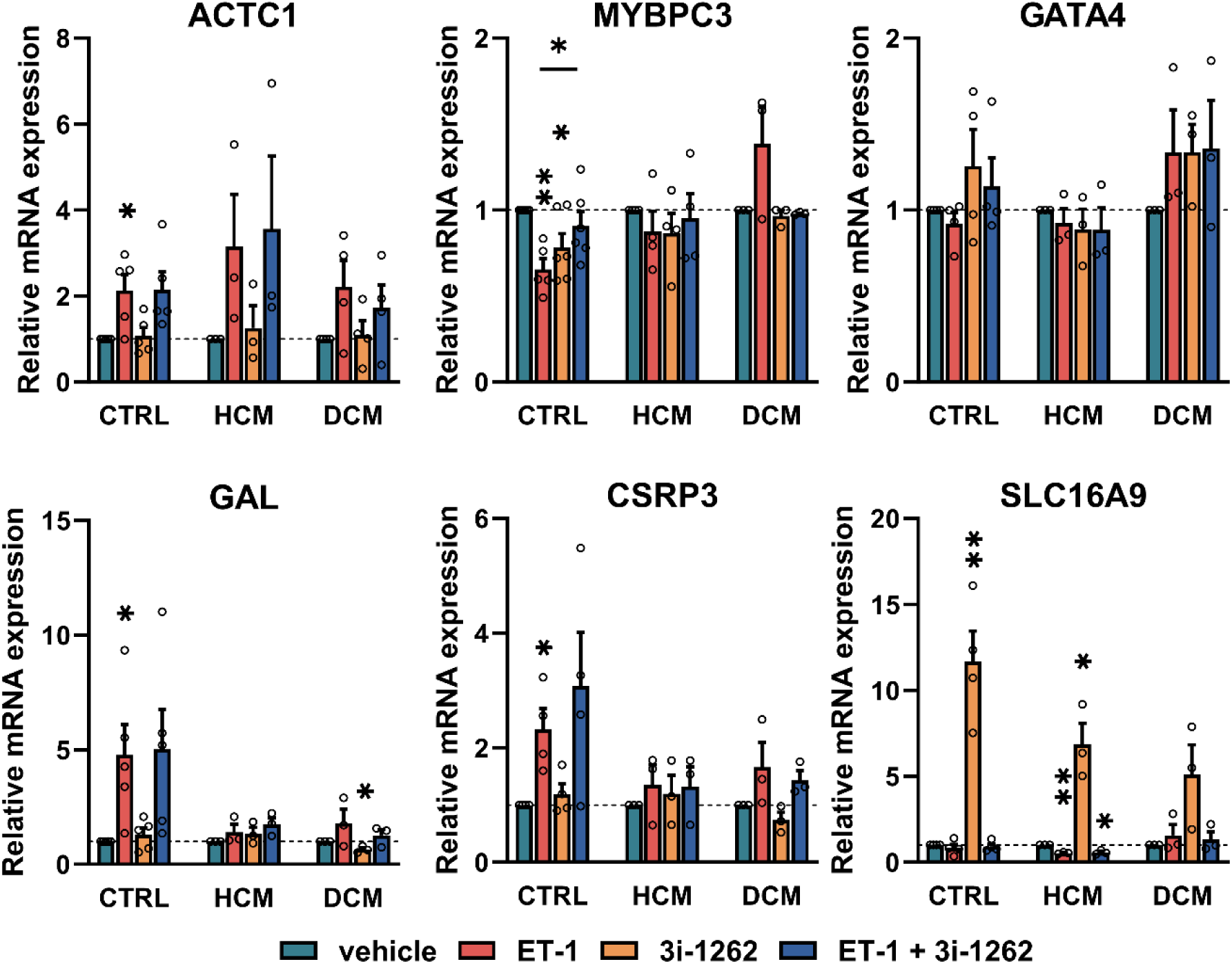
Gene expression of cardiomyocyte structural and regulatory genes in hiPSC-cardiomyocytes after 24 hours of endothelin-1 exposure and 3i-1262 treatment. Results are presented as mean + SEM and normalized to the respective vehicle sample. n = 3–6, *p<0.05; **p<0.01 compared to the respective vehicle sample, or as indicated (independent samples t-test or Welch’s t-test, as appropriate). CTRL, control cardiomyocytes; HCM, HCM cardiomyocytes; DCM, DCM cardiomyocytes; ET-1, endothelin-1.

### Baseline differences in contraction kinetics in control, HCM and DCM engineered heart tissues

To investigate contraction kinetics in control and patient-derived cardiomyocytes, engineered heart tissues (EHTs) were generated ^14^. Over the 40-day period, control cardiomyocyte EHTs maintained their structural integrity and continued beating until the end of the experiment (Figure 4A). Two out of three HCM EHTs remained viable until day 40, whereas one fractured on day 38. Among the DCM EHTs, one had ceased beating by day 18, a second by day 22, and the third by day 40. Representative videos of the EHTs from each line are shown in the Supplementary videos 1–9.

**Figure 4.**
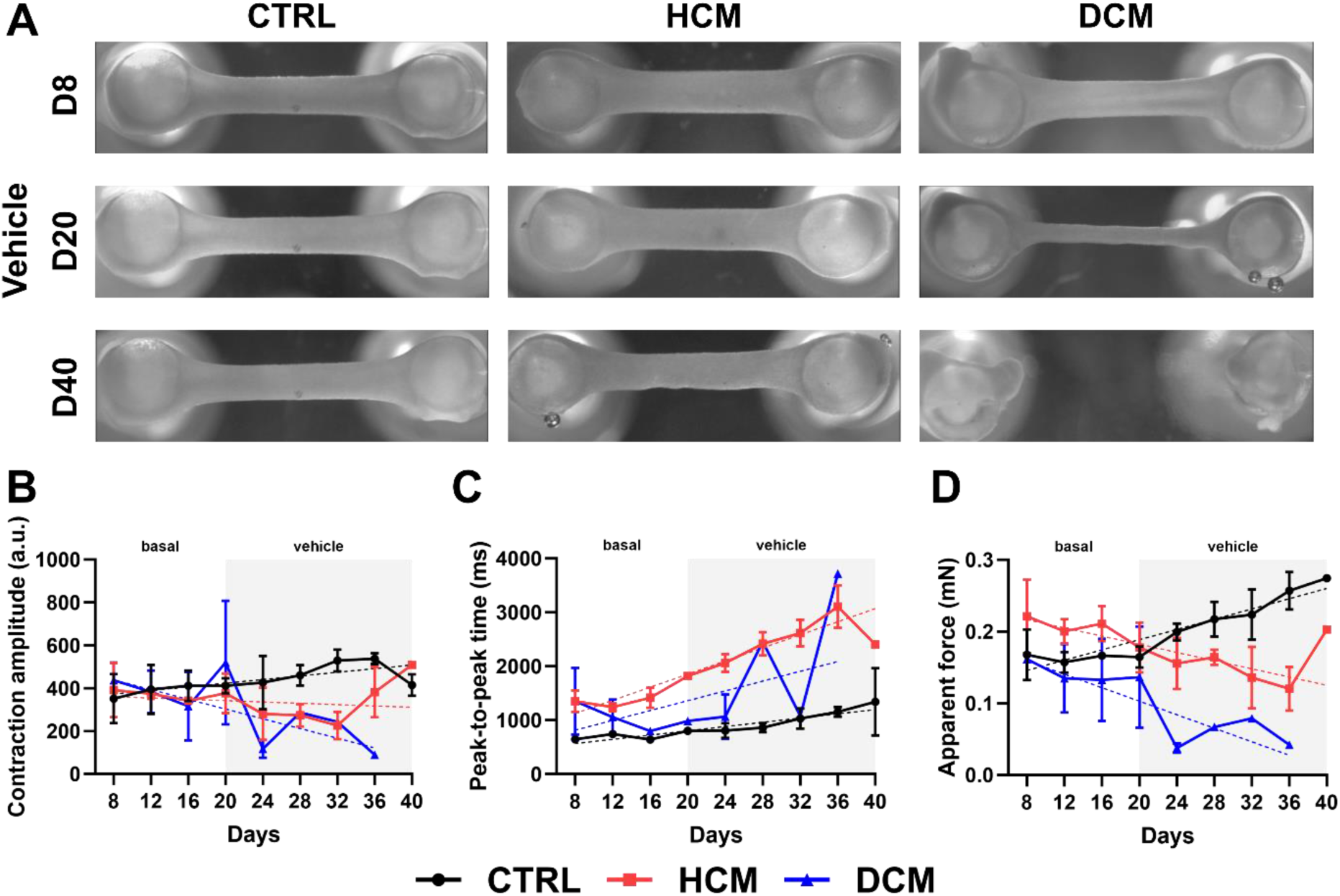
Contraction kinetics of engineered heart tissues (EHTs). A) Representative images of EHTs on days 8, 20, and 40 (D8, D20, D40, respectively). Basal contraction amplitude (a.u.) (B), peak-to-peak time (ms) (C), and apparent force (mN) (D) of EHTs. Results are presented as mean ±SEM. For control EHTs n = 3 throughout the study period, for HCM EHTs n = 3 on days 6 – 38 and n = 2 on day 40. For DCM EHTs, n = 3 from days 6– 16, n = 2 on days 18–30, and n = 1 from days 32–38 due to EHT fracture or cessation of beating. Data were analyzed using a linear mixed-effects model with group, day, and their interaction as fixed effects and EHT as a random intercept. Pairwise comparisons were Bonferroni-adjusted. The dashed line indicates the simple linear regression fit. CTRL, EHTs generated from control hiPSC-cardiomyocytes; HCM, EHTs generated from HCM patient-derived hiPSC-cardiomyocytes; DCM, EHTs generated from DCM patient-derived hiPSC-cardiomyocytes; a.u., arbitrary units.

HCM EHTs exhibited significantly lower contraction amplitude than control based on the estimated marginal means (p = 0.017), whereas DCM EHTs did not differ significantly from control (p = 0.236). However, over the 40-day period, linear mixed-effects models accounting for repeated measurements indicated that time was not a significant predictor of contraction amplitude in any comparison (HCM vs control: p = 0.779; DCM vs control: p = 0.811), indicating that contraction amplitude did not change significantly over time (Figure 4B). Notably, the model may not have fully captured both temporal changes and the lower contraction amplitude in DCM EHTs, likely due to the reduced number of EHTs at later time points rather than a true absence of effect.

Peak-to-peak time was significantly influenced by culture duration (p = 0.010), increasing across all groups and indicating a general slowing of contraction (Figure 4C). HCM EHTs exhibited a substantially higher estimated marginal mean of peak-to-peak time than control cells (p < 0.001). Despite this, the temporal trend was comparable between these groups, suggesting similar rates of change over time. Although the linear mixed-effects model did not detect a significant difference in peak-to-peak time between control and DCM EHTs, likely due to the limited number of viable EHTs, a significant difference in temporal trajectories was predicted (p = 0.028), with DCM EHTs exhibiting a faster increase in peak-to-peak time (Figure 4C). In force measurements, control, HCM and DCM EHTs followed distinct temporal trajectories with control cells showing an increase in apparent force over time, while HCM and DCM EHTs exhibited a decrease (p = 0.020 for HCM, p = 0.024 for DCM, compared to control) (Figure 4D). Additional contraction parameters, including relaxation time, contraction transient durations (90-90, 50-50, and 10-10 transients), and fractional shortening (%), are presented in Supplementary figure 5. These data demonstrate differences in transient durations between the DCM and control EHTs, as well as distinct trajectories of fractional shortening across both disease lines compared with control EHTs. Representative immunofluorescence images of EHT sections on day 40 are presented in Supplementary figure 6. Cardiac troponin T-positive tissue was observed in all groups, while visual inspection suggested a more peripheral distribution of DAPI staining in DCM EHTs; however, this was not quantitatively assessed.

### Endothelin-1 and 3i-1262-induced changes in the cardiomyocyte contraction kinetics

To assess the effects of endothelin-1 and compound 3i-1262 on contraction kinetics, cells were treated with endothelin-1, 3i-1262, or a combination of both starting from 20 days post-EHT generation. In the 3i-1262-treated groups, spontaneous beating ceased at variable time points across the different lines: in the control group, two EHTs stopped beating on day 20, when treatment was initiated, while the third ceased on day 24. Among the HCM EHTs, cessation occurred on days 22, 24, and 38. In the DCM EHTs, one EHT had already stopped beating by day 16, prior to starting 3i-1262 treatment, another stopped beating on day 20, and the third on day 24. Consequently, contraction amplitude, peak-to-peak interval, and apparent force could no longer be quantified after beating cessation (Supplementary video 10).

In control EHTs, contraction amplitude and peak-to-peak time were not significantly affected by endothelin-1 treatment,Figure 5 and the temporal trajectories did not differ between vehicle and endothelin-1 conditions (Figure 5A – B). Apparent force increased over time (p < 0.001), indicating a gradual strengthening of contractions, and endothelin-1 treatment did not alter this trajectory (Figure 5C). In untreated HCM EHTs, contraction amplitude decreased over time (p = 0.004); however, endothelin-1 treatment significantly increased contraction amplitude (p < 0.001) (Figure 5D). Furthermore, the temporal trajectory differed between treatments, suggesting that endothelin-1 treatment modulated the time-dependent changes in contraction amplitude. In contrast, peak-to-peak time was not significantly affected by time or treatment, and its temporal trajectory remained similar across treatment conditions (Figure 5E). Apparent force decreased over time in vehicle-treated HCM EHTs (p = 0.023), whereas endothelin-1 significantly modified its temporal trajectory (p < 0.001), shifting it from a declining trend to an increasing trend (Figure 5F).

**Figure 5.**
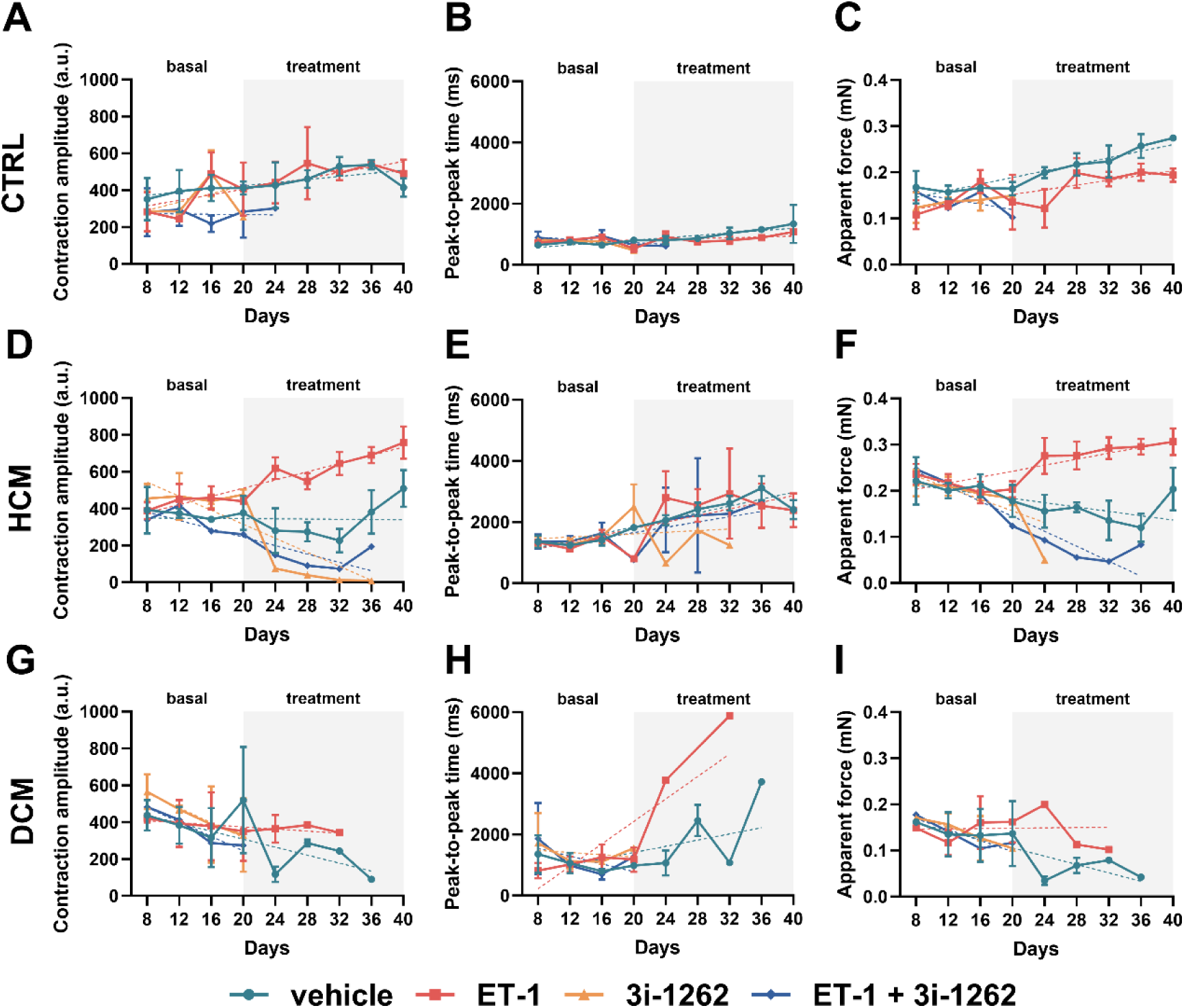
The effect of endothelin-1 and 3i-1262 treatments on contraction kinetics of engineered heart tissues (EHTs). Contraction amplitude (a.u.), peak-to-peak time (ms), and apparent force (mN) for control (A–C), HCM (D–F), and DCM (G–I) EHTs. Results are presented as mean ±SEM. The exact number of replicates (n) for each analysis is indicated in Supplementary tables 2 – 4. Data were analyzed using a linear mixed-effects model with group, day, and their interaction as fixed effects and EHT as a random intercept. Pairwise comparisons were Bonferroni-adjusted. The dashed line indicates the simple linear regression fit. CTRL, EHTs generated from control hiPSC-cardiomyocytes; HCM, EHTs generated from HCM patient-derived hiPSC-cardiomyocytes; DCM, EHTs generated from DCM patient-derived hiPSC-cardiomyocytes; ET-1, endothelin-1; a.u., arbitrary units.

In DCM EHTs, contraction amplitude did not change significantly over time and was not affected by the endothelin-1 treatment, with similar temporal trajectories observed across treatment conditions (Figure 5G). Peak-to-peak time did not change significantly over time and was not affected by treatment; however, its temporal trajectory differed between treatments (p < 0.001), predicting that endothelin-1 induced a more pronounced slowing of beating than vehicle treatment in DCM EHTs (Figure 5H). Finally, apparent force was significantly influenced by endothelin-1 treatment (p = 0.002); however, the temporal trajectory was not affected (Figure 5I). Additional contraction parameters following endothelin-1 and 3i-1262 treatments are shown in Supplementary figure 7. Endothelin-1 treatment did not alter these parameters in control EHTs; however, it modified the trajectory of fractional shortening in both DCM and HCM lines from a declining to an increasing pattern. In addition, it increased the 10-10 transient in both disease lines, which may indicate a prolonged relaxation phase. Representative immunofluorescence images of endothelin-1-treated EHT sections are presented in Supplementary figure 8, showing cardiac troponin T-positive tissue and a similarly peripheral distribution of DAPI staining in DCM EHTs as observed in vehicle-treated EHTs.

### Functional analysis of EHTs after β-adrenergic stimulation with isoprenaline

On day 40, we investigated the effects of β-adrenergic stimulation on contractile function of EHTs by exposing the tissues to 1 µM isoprenaline and recording their responses at 5, 10, 30, and 60 minutes. DCM EHTs were excluded from the endpoint analysis due to either fracture or cessation of beating. In control EHTs, contraction amplitude did not change following isoprenaline exposure, regardless of prior treatment with vehicle or endothelin-1 during the preceding 40-day period (p = 0.725; Figure 6A). Peak-to-peak time decreased as expected (p = 0.005; Figure 6B), reflecting increased contraction frequency upon β-adrenergic stimulation, and the response was similar in both vehicle and ET-1 groups. The apparent contraction force was higher in vehicle-treated than endothelin-1-treated tissues (p = 0.003; Figure 6C), consistent with observations during the preceding 40-day treatment period. The temporal trajectories of apparent force diverged (p = 0.032), decreasing in the vehicle group while remaining stable in the endothelin-1 group, suggesting that prior endothelin-1 exposure may modulate the apparent force response to β-adrenergic stimulation.

**Figure 6.**
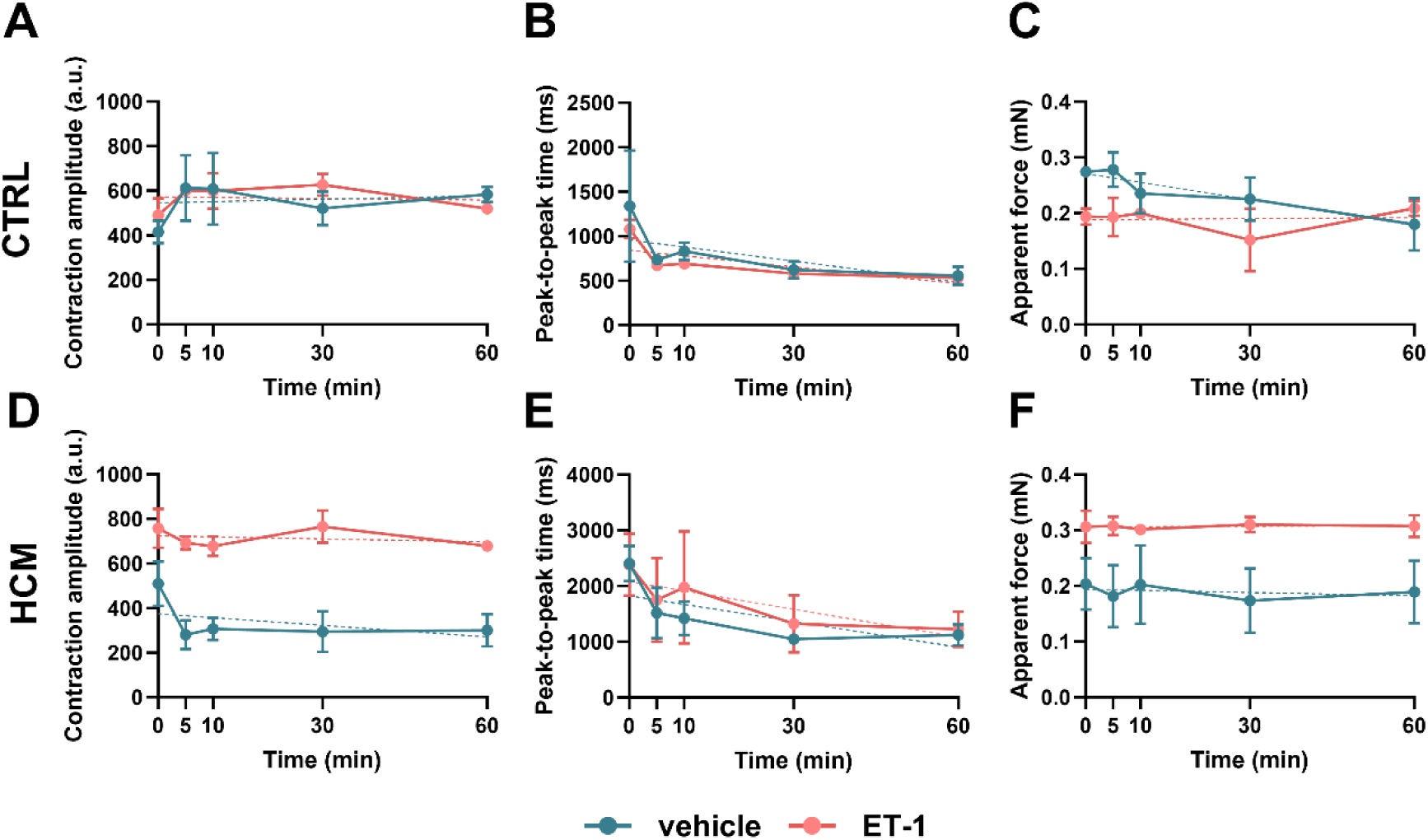
Contraction kinetics of hiPSC-derived engineered heart tissues in response to β-adrenergic stimulation with isoprenaline. Contraction amplitude (a.u.), peak-to-peak time (ms), and apparent force (mN) for control EHTs in panels A–C and HCM EHTs in panels D–F. Results are presented as mean ±SEM. Data were analyzed using a linear mixed-effects model with group, day, and their interaction as fixed effects and EHT as a random intercept. Pairwise comparisons were Bonferroni-adjusted. For the vehicle group, control n = 3 on days 6–40; HCM n = 3 on days 6–38 and n = 2 on day 40. For ET-1-treated EHTs, control and HCM n = 3 on days 6–40. The dashed line indicates the simple linear regression fit. CTRL, EHTs generated from control hiPSC-cardiomyocytes; HCM, EHTs generated from HCM patient-derived hiPSC-cardiomyocytes; ET-1, endothelin-1; a.u., arbitrary units.

In HCM EHTs, contraction amplitude was higher in tissues treated with endothelin-1 than in vehicle controls, as observed over the 40-day treatment period (p<0.001), and remained stable throughout the 60-minute isoprenaline exposure (Figure 6D). Unexpectedly, peak-to-peak time was unaffected by isoprenaline in both treatment groups (Figure 6E). Apparent force was elevated in endothelin-1-treated tissues and remained stable over the measurement period, with similar temporal trajectories between the groups (Figure 6F).

## Discussion

In this study, we modeled HCM and DCM using patient-specific hiPSC lines harboring *MYBPC3* (Gln1061X) and *LMNA* (S143P) mutations, respectively. Our findings provide important insights into the transcriptional, protein-level and functional alterations associated with HCM and DCM caused by these mutations, highlighting mutation-specific disease mechanisms with potential relevance for therapeutic development.

Endothelin-1 is widely used as a pro-hypertrophic stimulus in cardiac research and has been shown to induce hypertrophic remodeling in cardiomyocytes, including changes in hypertrophic marker gene expression ^19,35–40^. In the present study, control cardiomyocytes exhibited a hypertrophic response to endothelin-1 after 24 hours of exposure, including upregulation of *NPPB*, *NPPA*, and *ACTA1*, downregulation of *MYH6*, and increased proBNP protein levels. In contrast, cardiomyopathy patient-derived cardiomyocytes exhibited an attenuated response to endothelin-1, suggesting altered sensitivity to the stimulus. Consistent with this, previous studies have reported pronounced endothelin-1-induced phenotypic remodeling in HCM hiPSC-derived cardiomyocytes, including altered gene expression, increased cell size, and myofibrillar disarray ^39^. Similarly, we have previously demonstrated genotype-dependent transcriptional responses to mechanical stretch ^30^. Together, these findings support the concept that cardiomyocyte stress responses are shaped by both genetic background and the applied stimulus, resulting in context-dependent remodeling programs.

To investigate functional phenotypes in a 3-dimensional cardiac model, we generated EHTs from control and cardiomyopathy patient-derived hiPSC-cardiomyocytes. While functional EHTs were successfully generated from control, DCM, and HCM cardiomyocytes, the DCM patient-derived EHTs exhibited altered morphology and progressively lost structural integrity during maintenance, ultimately fracturing or ceasing to beat. To our knowledge, this is the first report describing such pronounced structural instability of *LMNA*-mutated hiPSC-cardiomyocyte-derived DCM EHTs under these conditions.

The DCM cardiomyocytes used in the study carry a serine-to-proline substitution at position p.S143P, introducing a structurally disruptive kink within the α-helical rod domain near the linker region^41^ . This alteration is predicted to perturb protein-protein interactions involving residue p.146, which are essential for the lateral assembly of lamin filaments and the formation of a stable, organized nuclear lamina. We speculate that this nuclear structural defect compromises the ability of DCM cardiomyocytes to withstand the mechanical load imposed by the three-dimensional EHT environment, consistent with the hypothesis that mutant lamins increase nuclear fragility, particularly in mechanically stressed tissues ^42^.

Although the fracture and cessation of beating observed in our DCM EHT model may support this hypothesis, the phenotype is unlikely to represent a general consequence of lamin dysfunction, as cardiomyocytes carrying other *LMNA* variants, including p.H222P^43^, c.357-2A>G^44^, and p.L204P^45^, have been successfully incorporated into EHTs. Notably, these variants may exert distinct structural and mechanistic effects from p.S143P. Located in the coil 1B region of the lamin rod domain, the p.S143P substitution may be particularly disruptive to lamin filament assembly and potentially to nucleo-cytoskeletal coupling through the LINC complex, thereby rendering cardiomyocytes particularly vulnerable to the increased mechanical demands of the three-dimensional EHT environment^41^. Thus, the limited structural stability of the DCM EHTs in the present study may reflect variant-specific vulnerability rather than a general consequence of the *LMNA*-related defect. This interpretation is also consistent with studies showing successful generation of engineered cardiac tissues from cardiomyocytes carrying other DCM-causing mutations, including variants in leiomodin 2 (*LMOD2*)^46^, titin (*TTN*)^47^, RNA binding motif protein 20 (*RBM20*)^17^, phospholamban (*PLN*)^48^, and cardiac troponin T (*TNNT2*)^49^. Collectively, these findings suggest that EHT fracture and cessation of beating in the present model are unlikely to result simply from the presence of a pathogenic cardiomyopathy mutation. It is therefore tempting to speculate that they may instead arise from mutation-specific structural vulnerability associated with the p.S143P *LMNA* variant. Interestingly, irrespective of the causative mutation or the gene involved, force generation in DCM EHTs has consistently been reported to decline over time^17,46,47^ in agreement with our data, where linear mixed-effects modeling suggested a progressive decline in apparent force in DCM EHTs.

While the HCM EHTs remained structurally intact, we observed reduced force generation in HCM EHTs, consistent with previous genetic HCM models reporting impaired contractile force in *MYBPC3*-associated and other sarcomeric mutation backgrounds ^50–53^. However, alterations in force generation are not uniform across studies, as several HCM models have also demonstrated increased force generation ^53–59^. In addition, Weber et al. (2025) reported substantial cell-to-cell variability in force generation in *MYH7*-mutant hiPSC-cardiomyocytes, described as “contractile imbalance”, further highlighting the heterogeneous and mutation-dependent nature of force alterations in HCM ^60^.

Several mechanisms have been proposed to explain the altered force generation observed in HCM cardiomyocytes. Truncating *MYBPC3* mutations and loss of cardiac myosin binding protein (cMyBP-C) have been suggested to dysregulate myosin head conformations by destabilizing the super-relaxed state (SRX) and shifting myosin heads towards the more active disordered relaxed state (DRX), thereby altering contractility and increasing energetic demand ^61–63^. More broadly, impaired force generation in HCM models has been attributed to defective sarcomere formation resulting in reduced myofibrillar density, perturbed cross-bridge cycling, increased myofilament calcium sensitivity leading to diastolic dysfunction, and metabolic disturbances ^50^. Collectively, these findings suggest that altered force generation in HCM is likely multifactorial, reflecting both primary mutation-specific effects and secondary cellular adaptations.

In addition to its pro-hypertrophic effects, endothelin-1 is recognized as one of the most potent endogenous positive inotropic mediators in the heart ^37,64^. Surprisingly, endothelin-1 did not significantly affect contractile parameters in control EHTs, whereas it increased contraction amplitude and apparent force generation in HCM EHTs compared with vehicle-treated controls. Consistent with this, *MYBPC3*-mutant hiPSC-derived cardiomyocytes have previously been reported to exhibit altered inotropic effect after a 7-day exposure to endothelin-1 and other agents ^52^. Together, these results suggest that hiPSC-derived cardiomyocytes display genotype-specific responses to endothelin-1, reflected in differential effects at both the gene expression and contractile function levels.

Hypertrophic and dilated cardiomyopathies remain incurable disorders, and current treatments are primarily focused on symptom management, although disease-modifying therapies are emerging ^2^. However, targeting transcription factor networks offers a promising avenue for disease modulation, for example, in pathological cardiac hypertrophy ^65–67^. The GATA4-targeted compound 3i-1262 has previously been shown to reduce endothelin-1-induced proBNP and *NPPB* expression, as well as stretch-induced *NPPB* expression, in hiPSC-derived cardiomyocytes ^19,28,30^. However, in the present study, we did not observe comparable effects on hypertrophic marker expression in control or patient-derived cardiomyocytes. In addition to assessing gene and protein expression, we investigated the effects of 3i-1262 on cardiomyocyte function, hypothesizing that modulation of transcription factor networks might improve contractile function. Unexpectedly, treatment with 3i-1262 resulted in cessation of beating in all EHTs. This finding highlights the challenges associated with pharmacological modulation of transcription factor networks. As transcription factors regulate gene programs essential for both normal cardiomyocyte function and pathological remodeling, their pharmacological modulation requires precise control ^67^.

We acknowledge several limitations of this study. First, hiPSC-derived cardiomyocytes do not fully recapitulate the phenotype of adult cardiomyocytes because they retain an immature phenotype ^68,69^, which may influence hypertrophic responses and contractile function within the EHT model. Second, the study exclusively utilized hiPSC-derived cardiomyocytes and therefore did not capture the multicellular interactions present in native cardiac tissue ^70^. Finally, only one patient-derived hiPSC line was included for each disease and compared with a single control line, and no isogenic controls were available. Consequently, the findings may not fully reflect the heterogeneity of HCM and DCM, and further experiments in additional patient and isogenic lines will be required to confirm the generalizability of these findings.

In conclusion, this study demonstrates that hiPSC-cardiomyocytes derived from patients with genetic cardiomyopathy exhibit distinct expression patterns of hypertrophy-associated genes and proBNP protein as well as functional characteristics compared with control cardiomyocytes. Notably, EHTs generated from DCM patient-derived hiPSC-cardiomyocytes harboring the *LMNA* S143P mutation failed to maintain structural integrity. In addition, hiPSC-cardiomyocytes displayed genotype-specific responses to hypertrophic stimulation and pharmacological treatment. Together, these findings highlight the utility of patient-derived hiPSC-based models for investigating disease mechanisms and pharmacological responses in inherited cardiomyopathies.

## Acknowledgements

The facilities and expertise of the BI Histology core facility at the University of Helsinki, supported by HiLIFE, are gratefully acknowledged. We thank Dr Lotta Pohjolainen, Annika Korvenpää and Tatiana Tarkhova for expert technical assistance.

## Supplementary materials list

The supplementary materials include Supplementary tables, Supplementary figures, and Supplementary videos, along with their respective legends.

## Supplementary video legends

Supplementary Video 1. Representative video of an engineered heart tissue (EHT) generated from control hiPSC-cardiomyocytes on day 8.

Supplementary Video 2. Representative video of an EHT generated from control hiPSC-cardiomyocytes on day 20, recorded 30 minutes after the vehicle treatment.

Supplementary Video 3. Representative video of an EHT generated from control hiPSC-cardiomyocytes on day 40, recorded 30 minutes after the vehicle treatment.

Supplementary Video 4. Representative video of an EHT generated from hypertrophic cardiomyopathy (HCM) patient-derived hiPSC-cardiomyocytes on day 8.

Supplementary Video 5. Representative video of an EHT generated from HCM patient-derived hiPSC-cardiomyocytes on day 20, recorded 30 minutes after the vehicle treatment.

Supplementary Video 6. Representative video of an EHT generated from HCM patient-derived hiPSC-cardiomyocytes on day 40, recorded 30 minutes after the vehicle treatment.

Supplementary Video 7. Representative video of an EHT generated from dilated cardiomyopathy (DCM) patient-derived hiPSC-cardiomyocytes on day 8.

Supplementary Video 8. Representative video of an EHT generated from DCM patient-derived hiPSC-cardiomyocytes on day 20, recorded 30 minutes after the vehicle treatment.

Supplementary Video 9. Representative video of an EHT generated from DCM patient-derived hiPSC-cardiomyocytes on day 40, recorded 30 minutes after the vehicle treatment.

Supplementary Video 10. Representative video of an EHT generated from control hiPSC-cardiomyocytes on day 20, recorded 30 minutes after treatment with 30 µM 3i-1262. For the respective vehicle-treated EHTs, refer to Supplementary Video 2.

---

**Supplementary figure 1.**
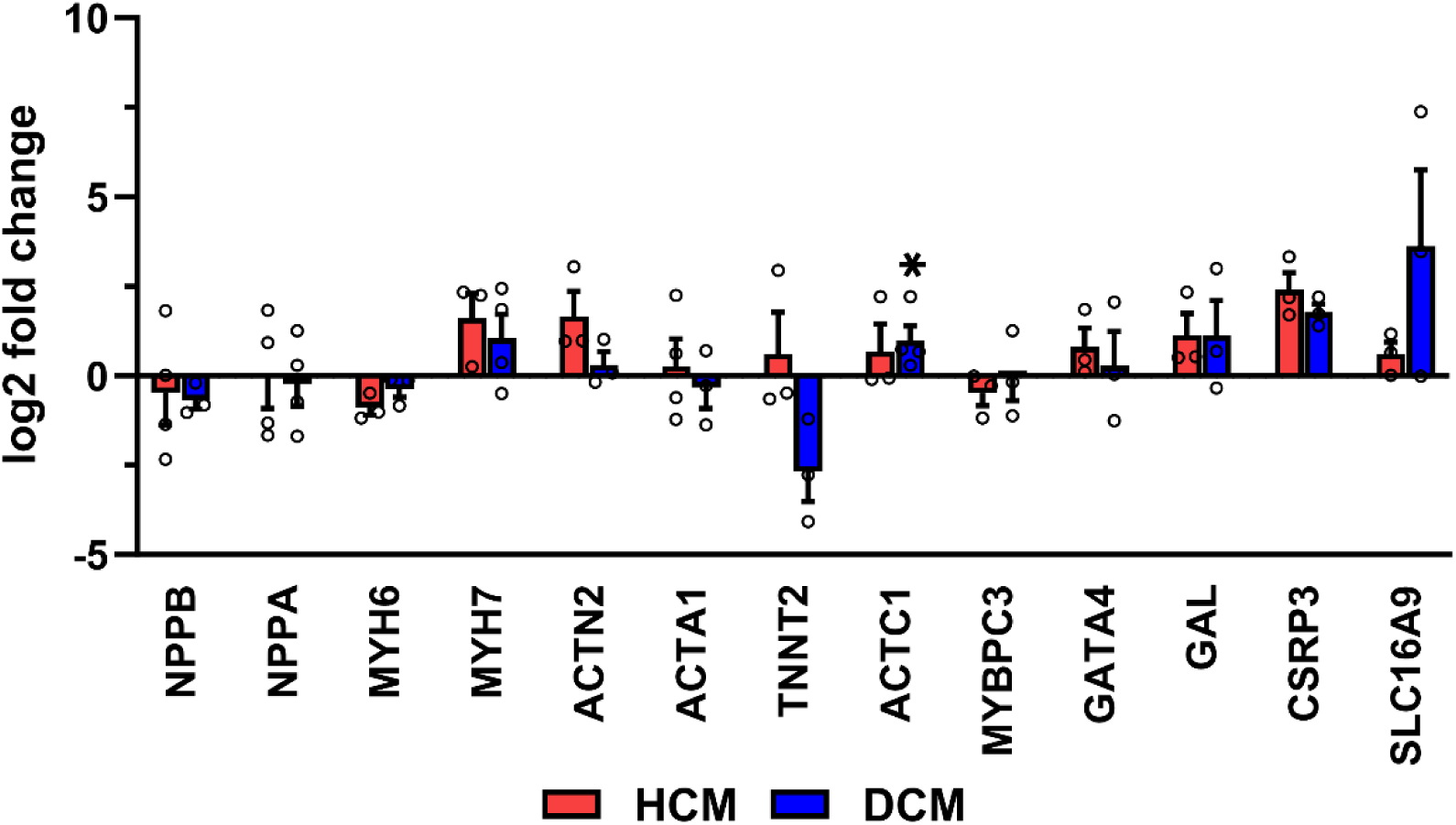
Basal gene expression in HCM and DCM hiPSC-cardiomyocytes. Data are presented as mean ±SEM (n = 3 – 4). *p<0.05, compared to the control cardiomyocytes (independent samples t-test or Welch’s t-test, as appropriate). HCM, HCM patient-derived hiPSC-cardiomyocytes; DCM, DCM patient-derived hiPSC-cardiomyocytes.

**Supplementary figure 2.**
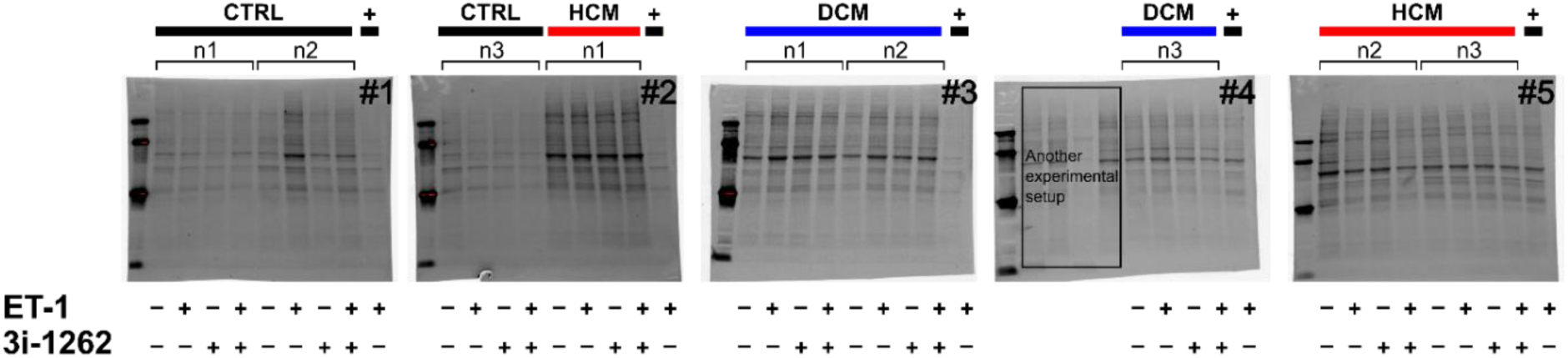
Total protein levels of hiPSC-cardiomyocytes at baseline and after endothelin-1 and 3i-1262 treatment. CTRL, control hiPSC-cardiomyocytes; HCM, HCM patient-derived hiPSC-cardiomyocytes; DCM, DCM patient-derived hiPSC-cardiomyocytes; ET-1, endothelin-1. The + sign indicates the positive control: control hiPSC-cardiomyocytes from an independent differentiation that were treated with endothelin-1 for 24 hours.

**Supplementary figure 3.**
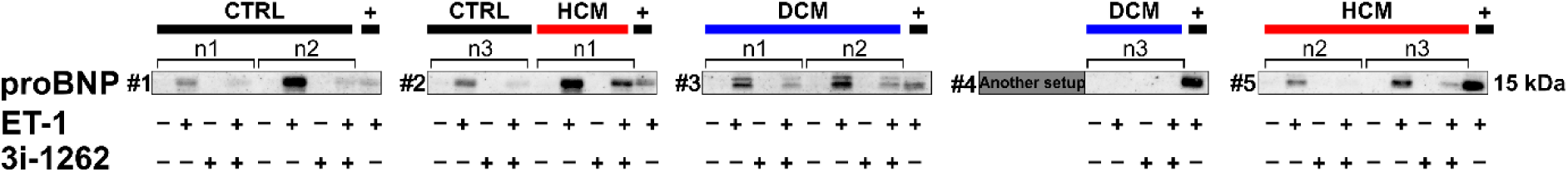
Representative Western blots of pro-B-type natriuretic peptide (proBNP; 15 kDa) in hiPSC-cardiomyocytes under basal conditions and treatment with endothelin-1 and/or 3i-1262. Corresponding total protein membranes are provided in Supplementary figure 2, with membranes labeled according to the same numbering scheme as in this figure (e.g., membrane #1 corresponds to total protein #1). CTRL, control cardiomyocytes; HCM, HCM patient-derived hiPSC-cardiomyocytes; DCM, DCM patient-derived hiPSC-cardiomyocytes; ET-1, endothelin-1. The + sign indicates the positive control: control hiPSC-cardiomyocytes from an independent differentiation that were treated with endothelin-1 for 24 hours.

**Supplementary figure 4.**
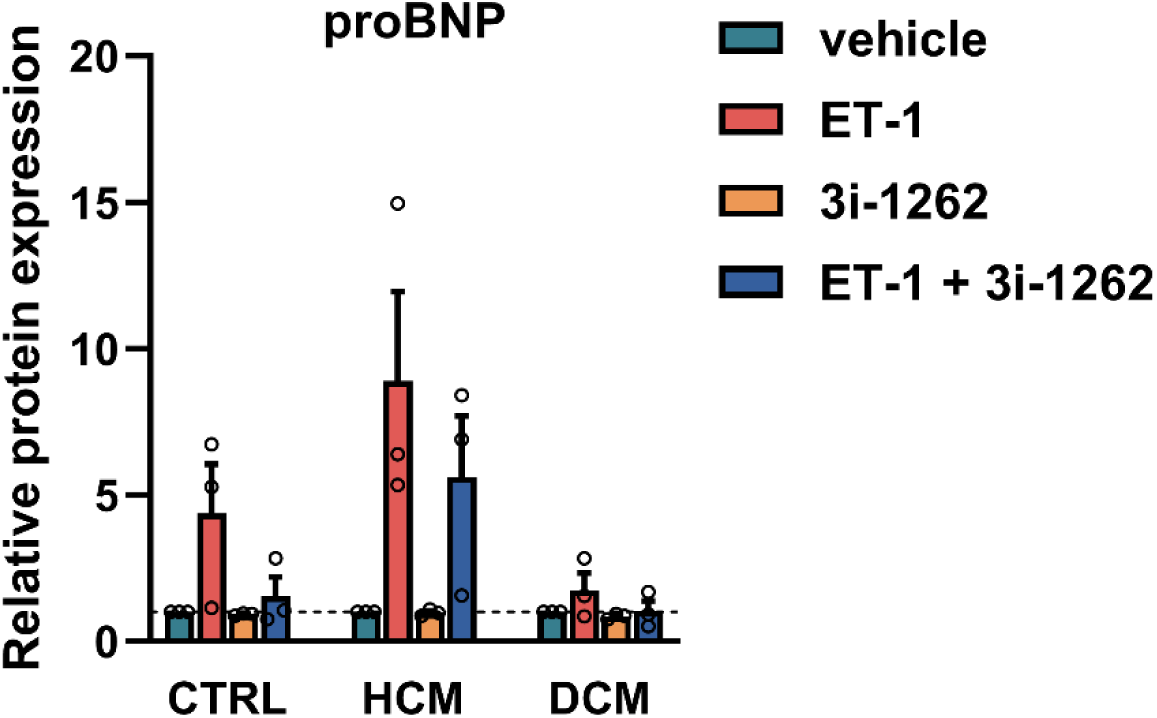
Expression of proBNP in hiPSC-cardiomyocytes. Results are presented as mean + SEM and normalized to the respective vehicle sample. n = 3, independent samples t-test or Welch’s t-test, as appropriate. CTRL, control cardiomyocytes; HCM, HCM cardiomyocytes; DCM, DCM cardiomyocytes; ET-1 = endothelin-1. Original total protein images and immunoblots are provided in Supplementary figure 2 and Supplementary figure 3.

**Supplementary figure 5.**
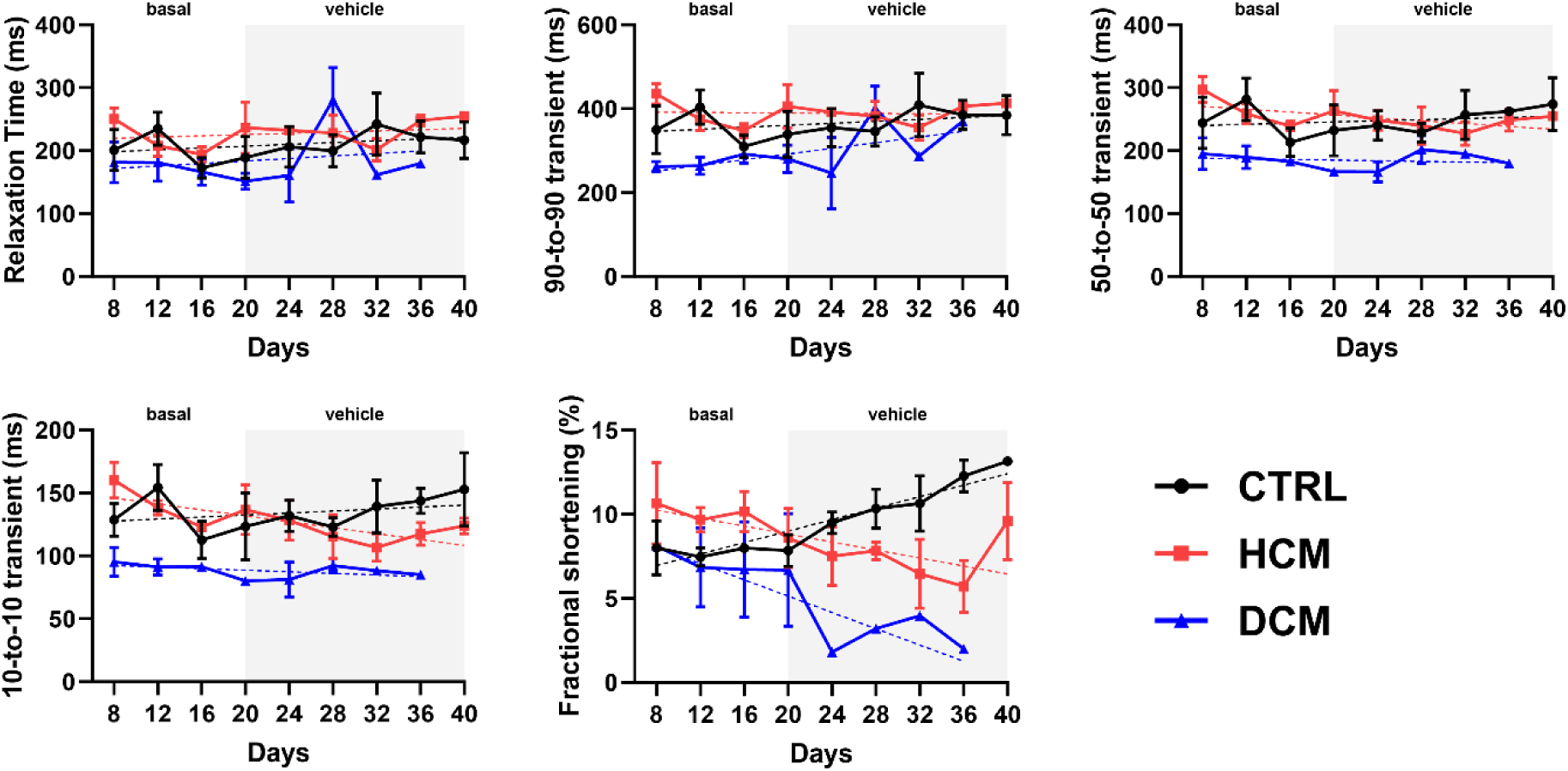
Contraction kinetics of engineered heart tissues (EHTs). Data are presented as mean ±SEM. For EHTs generated using control hiPSC-cardiomyocytes, n = 3 throughout the study period. For EHTs generated using HCM patient-derived hiPSC-cardiomyocytes, n = 3 on days 6–38 and n = 2 on day 40. For DCM EHTs, n = 3 from days 6–16, n = 2 on days 18–30, and n = 1 from days 32–38 due to EHT fracture or cessation of beating. The dashed line represents the simple linear regression fit. CTRL, EHTs generated from control hiPSC-cardiomyocytes; HCM, EHTs generated from HCM patient-derived hiPSC-cardiomyocytes; DCM, EHTs generated from DCM patient-derived hiPSC-cardiomyocytes.

**Supplementary figure 6.**
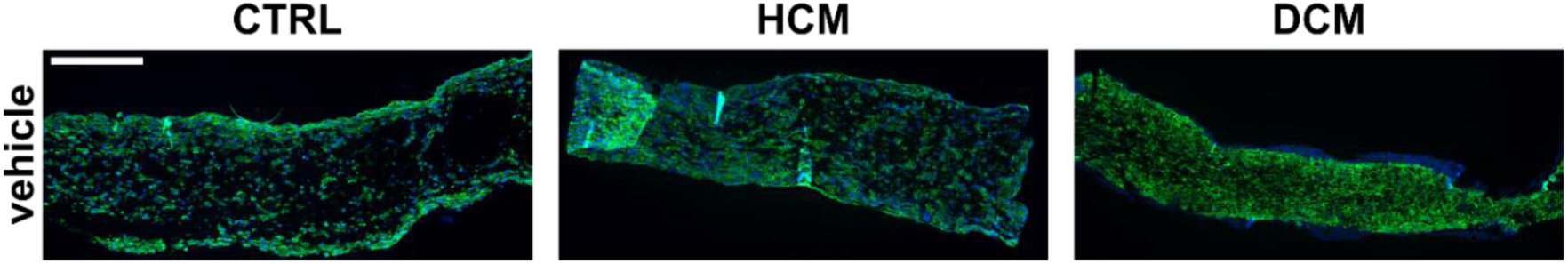
Immunofluorescence staining of engineered heart tissue (EHT) sections on day 40. hiPSC-cardiomyocyte EHTs were sectioned into 4 µM sections and stained for DNA (DAPI; blue) and cardiac troponin T (green). CTRL, EHTs generated from control hiPSC-cardiomyocytes; HCM, EHTs generated from HCM patient-derived hiPSC-cardiomyocytes; DCM, EHTs generated from DCM patient-derived hiPSC-cardiomyocytes. Scale bar: 500 µM.

**Supplementary figure 7.**
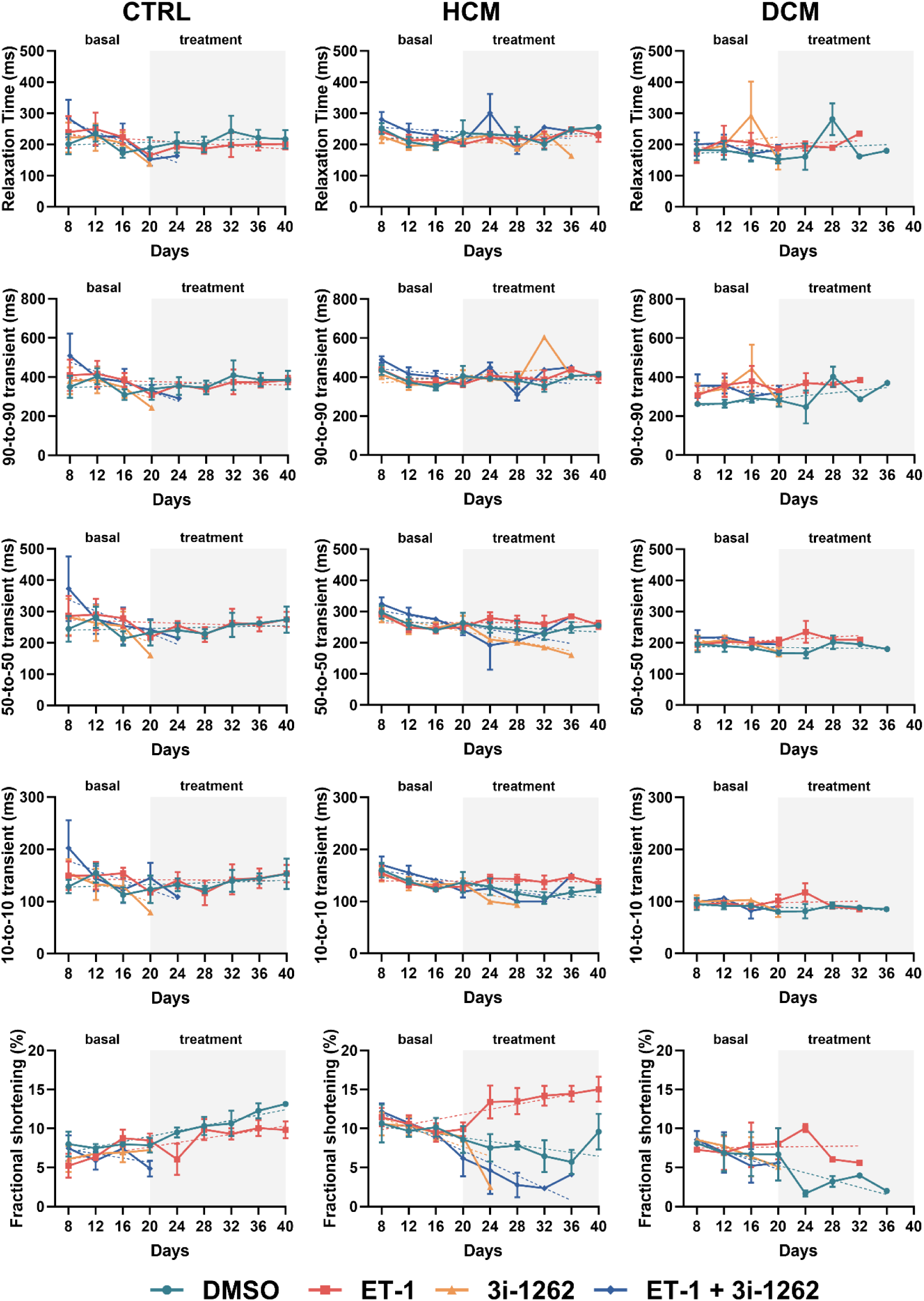
Effects of vehicle, endothelin-1, and 3i-1262 treatments on the contraction kinetics of engineered heart tissues (EHTs). Data are presented as mean ±SEM. The exact number of replicates (n) for each analysis are indicated in Supplementary tables 2 – 4. The dashed line represents the simple linear regression fit. CTRL, EHTs generated from control hiPSC-cardiomyocytes; HCM, EHTs generated from HCM patient-derived hiPSC-cardiomyocytes; DCM, EHTs generated from DCM patient-derived hiPSC-cardiomyocytes; ET-1, endothelin-1.

**Supplementary figure 8.**
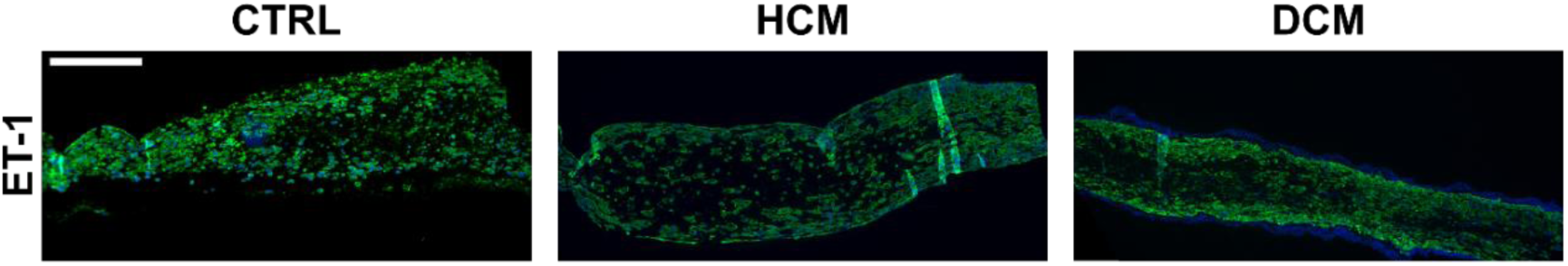
Immunofluorescence staining of endothelin-1-treated engineered heart tissue (EHT) sections on day 40. hiPSC-cardiomyocyte EHTs were sectioned into 4 µM sections and stained for DNA (DAPI; blue) and cardiac troponin T (green). CTRL, EHTs generated from control hiPSC-cardiomyocytes; HCM, EHTs generated from HCM patient-derived hiPSC-cardiomyocytes; DCM, EHTs generated from DCM patient-derived hiPSC-cardiomyocytes.; ET-1, endothelin-1. Scale bar: 500 µM.

**Supplementary table 1.** TaqMan assays used in qPCR experiments.

| Gene | Function | TaqMan® Gene Expression Assay ID |
| --- | --- | --- |
| *ACTB* | Housekeeping gene | 4352935E |
| *18S rRNA* | Housekeeping gene | 4352930E |
| *NPPB* | Hypertrophy marker | Hs01057466_g1 |
| *NPPA* | Hypertrophy marker | Hs00383230_g1 |
| *MYH6* | Sarcomeric gene | Hs01101425_m1 |
| *MYH7* | Sarcomeric gene | Hs01110632_m1 |
| *ACTC1* | Sarcomeric gene | Hs01109515_m1 |
| *TNNT2* | Sarcomeric gene | Hs00943911_m1 |
| *MYBPC3* | Sarcomeric gene | Hs00165232_m1 |
| *ACTN2* | Sarcomeric gene | Hs00153809_m1 |
| *ACTA1* | Sarcomeric gene | Hs00559403_m1 |
| *CSRP3* | Sarcomeric gene | Hs00185787_m1 |
| *GATA4* | Cardiac transcription factor | Hs00171403_m1 |
| *GAL* | Metabolic-related gene | Hs00544355_m1 |
| *SLC16A9* | Metabolic-related gene | Hs00415854_m1 |

**Supplementary table 2.** Number of EHTs generated using control hiPSC-cardiomyocytes (n) included throughout the study. The table presents the number of replicates for the treatment groups at different time points (days 6 – 40). The number of replicates varied due to EHT fracture or cessation of beating. N/A indicates that no data was available because measurements could not be performed due to EHT fracture or cessation of beating.

| Day | Vehicle | ET-1 | 3i-1262 | ET-1 + 3i-1262 |
| --- | --- | --- | --- | --- |
| 6 | 3 | 3 | 3 | 3 |
| 8 | 3 | 3 | 3 | 3 |
| 10 | 3 | 3 | 3 | 3 |
| 12 | 3 | 3 | 3 | 3 |
| 14 | 3 | 3 | 3 | 3 |
| 16 | 3 | 3 | 3 | 3 |
| 18 | 3 | 3 | 3 | 3 |
| 20 | 3 | 3 | 3 | 3 |
| 22 | 3 | 3 | 1 | 2 |
| 24 | 3 | 3 | 1 | 1 |
| 26 | 3 | 3 | N/A | N/A |
| 28 | 3 | 3 | N/A | N/A |
| 30 | 3 | 3 | N/A | N/A |
| 32 | 3 | 3 | N/A | N/A |
| 34 | 3 | 3 | N/A | N/A |
| 36 | 3 | 3 | N/A | N/A |
| 38 | 3 | 3 | N/A | N/A |
| 40 | 3 | 3 | N/A | N/A |

**Supplementary table 3.**
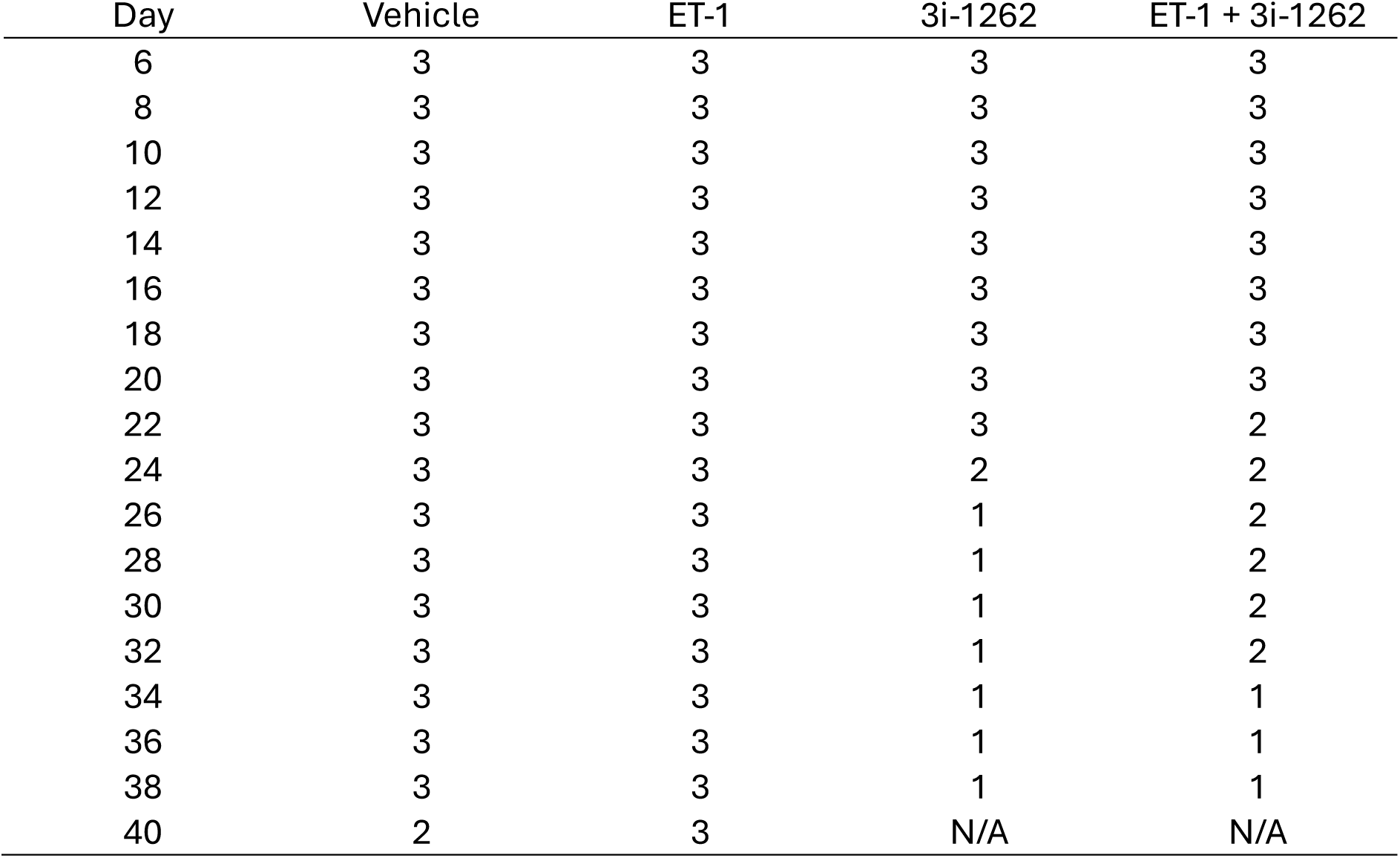
Number of EHTs generated using HCM hiPSC-cardiomyocytes (n) included throughout the study. The table presents the number of replicates for the treatment groups at different time points (days 6 – 40). The number of replicates varied due to EHT fracture or cessation of beating. N/A indicates that no data was available because measurements could not be performed due to EHT fracture or cessation of beating.

**Supplementary table 4.** Number of EHTs generated using DCM hiPSC-cardiomyocytes (n) included throughout the study. The table presents the number of replicates for the treatment groups at different time points (days 6 – 40). The number of replicates varied due to EHT fracture or cessation of beating. N/A indicates that no data was available because measurements could not be performed due to EHT fracture or cessation of beating.

| Day | Vehicle | ET-1 | 3i-1262 | ET-1 + 3i-1262 |
| --- | --- | --- | --- | --- |
| 6 | 3 | 3 | 3 | 3 |
| 8 | 3 | 3 | 3 | 3 |
| 10 | 3 | 3 | 3 | 3 |
| 12 | 3 | 3 | 3 | 3 |
| 14 | 3 | 3 | 3 | 3 |
| 16 | 3 | 3 | 3 | 3 |
| 18 | 2 | 3 | 2 | 2 |
| 20 | 2 | 3 | 2 | 2 |
| 22 | 2 | 3 | 1 | 1 |
| 24 | 2 | 2 | 1 | N/A |
| 26 | 2 | 1 | N/A | N/A |
| 28 | 2 | 1 | N/A | N/A |
| 30 | 2 | 1 | N/A | N/A |
| 32 | 1 | 1 | N/A | N/A |
| 34 | 1 | 1 | N/A | N/A |
| 36 | 1 | 1 | N/A | N/A |
| 38 | 1 | 1 | N/A | N/A |
| 40 | N/A | N/A | N/A | N/A |

